# Compartmentalized Nanozyme Transduction Enables Regenerative Molecular Amplification for Multiplexed miRNA-Based Prostate Cancer Stratification

**DOI:** 10.64898/2026.09.29.755280

**Authors:** Ying Ma, Xiang-Lan He, Jia-Qi Du, Meng-Meng Pan, Ming Wang, Guan-Yu Qu, Ming Jiang, Li Xu, Xu Yu

## Abstract

Circulating microRNAs (miRNAs) have emerged as promising liquid biopsy markers; however, their clinical translation is restricted by low abundance, biological heterogeneity, and the complexity of multiplex profiling. Herein, we develop a **t**argeted **<u>c</u>**ycling-mediated **<u>d</u>**uplex-**<u>s</u>**pecific **<u>n</u>**uclease (**DSN**)-**<u>nano</u>**zyme signal amplification platform (**TC-DSN-Nano**) that establishes a compartmentalized catalytic transduction strategy for multiplexed miRNA analysis. By encapsulating gold nanozymes within liposomal nanoreactors, this platform spatially separates molecular recognition from catalytic signal generation, preserving nanozyme activity during sensor construction. Upon target miRNA recognition, DSN-mediated recycling induces regenerative release of nanozyme-loaded liposomes from magnetic substrates, converting individual miRNA molecules into amplified catalytic outputs without conventional nucleic acid amplification. The platform enables programmable profiling of six prostate cancer (PCa) associated miRNAs in serum samples. Furthermore, machine learning-assisted integration of multiplexed miRNA signatures enables accurate discrimination among healthy donors, nonmetastatic PCa patients, and metastatic PCa patients, achieving an overall classification accuracy of 90.5%. This work establishes a general compartmentalized nanozyme transduction framework that integrates molecular recycling, catalytic amplification, and computational analysis for liquid biopsy-based cancer stratification.

**TOC:** 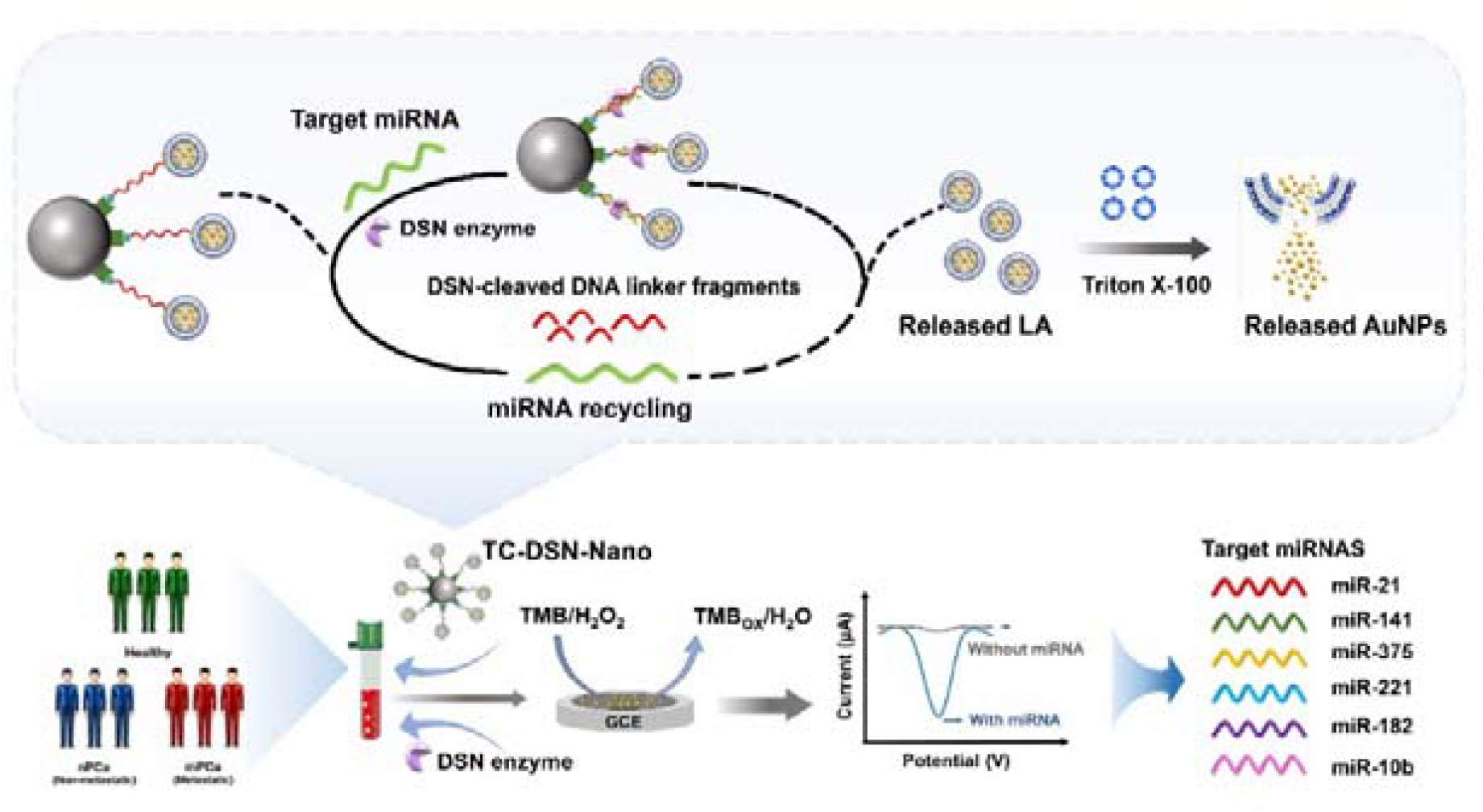

## 1. Introduction

Sensitive and multiplexed analysis of circulating nucleic acid biomarkers is increasingly important for liquid biopsy and molecular diagnostics. Among these biomarkers, circulating microRNAs (miRNAs), a class of evolutionarily conserved noncoding RNAs,^1-3^ have emerged as promising liquid biopsy biomarkers because their expression patterns are closely associated with cancer development and metastatic dissemination.^4-6^ Importantly, individual miRNAs often provide limited diagnostic information because of biological heterogeneity and interpatient variation, whereas multiplexed miRNA signatures can integrate complementary molecular information and improve disease classification.^7, 8^ Recent advances in multivariate statistical analysis and machine learning have further highlighted the value of combining multiple miRNA biomarkers to extract clinically relevant information that cannot be obtained from individual molecular markers alone.^9, 10^ Therefore, analytical platforms capable of quantitatively profiling multiple circulating miRNAs are highly desirable for translating miRNA signatures into clinically relevant molecular information.

However, accurate miRNA profiling in complex biological samples remains challenging because miRNAs are typically present at low abundance, comprise short sequences of only 18-24 nucleotides, and exhibit high sequence similarity.^11-13^ These characteristics make it difficult to distinguish closely related miRNAs while simultaneously generating sufficiently strong and quantitative signals from scarce target molecules. Although advanced technologies, including polymerase chain reaction (PCR),^14, 15^ next-generation sequencing,^16, 17^ and CRISPR-based assays,^18-20^ have substantially advanced miRNA analysis, their practical implementation can involve complex workflows, sophisticated probe designs, specialized instrumentation, or stringent reaction conditions.^12^ Thus, a key challenge is to develop a sensing strategy that can efficiently amplify the molecular recognition events of multiple low-abundance miRNAs while maintaining sequence-specific and quantitative signal transduction.

Nucleic acid amplification strategies have therefore been widely explored to enhance the analytical response of miRNA detection. Conventional approaches, including PCR,^21^ rolling circle amplification,^22, 23^ and hybridization-based cascade amplification,^24^ can provide substantial signal enhancement but often require complicated reaction schemes and are less straightforward to adapt to multiplexed molecular analysis. Alternatively, duplex-specific nuclease (DSN)-mediated target recycling provides an attractive strategy for regenerative amplification of molecular recognition events.^25-27^ DSN selectively cleaves the DNA strand within an RNA–DNA heteroduplex while preserving the RNA target, allowing the target miRNA to be regenerated and participate in successive rounds of hybridization and DNA cleavage. Thus, DSN can effectively multiply the number of molecular recognition events initiated by a single miRNA molecule. However, DSN-mediated recycling primarily amplifies molecular recognition events rather than directly generating amplified analytical signals, and additional transduction strategies mechanisms are required to efficiently convert these regenerated molecular events into detectable analytical outputs. Therefore, the integration of regenerative molecular amplification with catalytic signal amplification represents an attractive strategy for developing advanced miRNA sensing platforms.

Nanozymes are emerging as promising signal reporters because of their high catalytic activity, structural robustness, and compatibility with electrochemical readout.^28-30^ However, in many reported biosensing systems, nanozymes are directly functionalized with recognition probes or immobilized within sensing architectures, where surface modification and biomolecular adsorption can partially block catalytic active sites and reduce substrate accessibility.^31, 32^ Consequently, the intrinsic catalytic amplification capability of nanozymes can be compromised during sensor fabrication. Therefore, an ideal sensing architecture should not only integrate regenerative target recycling with nanozyme-based catalytic amplification but also spatially separate molecular recognition from catalytic signal generation to preserve nanozyme activity.^33-35^ To address this challenge, we introduced a liposomal compartment to spatially isolate the AuNP nanozymes from the recognition and transduction components, thereby protecting their catalytic surfaces during sensor construction while enabling controlled release upon membrane disruption. This compartmentalized design minimizes unnecessary interference with the nanozyme surface while allowing the released AuNPs to remain fully accessible for subsequent catalytic signal amplification.

Herein, we developed a **t**argeted **<u>c</u>**ycling-mediated **<u>DSN</u>**-**<u>nano</u>**zyme signal amplification platform (TC-DSN-Nano) that establishes a regenerative and compartmentalized nanozyme transduction strategy for multiplexed miRNA analysis. The system integrates magnetic bead-assisted DNA linkers for target-responsive DSN cleavage and regenerative miRNA recycling with liposome-compartmentalized AuNP nanozymes for spatially separated catalytic signal transduction. In this design, DSN-mediated target recycling amplifies molecular recognition events, while liposomal encapsulation protects the catalytic AuNPs during sensor construction and enables their controlled release for subsequent electrochemical signal amplification. Thus, the two amplification processes are functionally separated but mechanistically coupled: DSN provides regenerative molecular amplification, whereas the released AuNP nanozymes Importantly, the recognition sequence can be readily programmed by changing the DNA linker, allowing the same signal transduction architecture to be adapted to different miRNA targets without redesigning the catalytic module. As a proof-of-concept application, TC-DSN-Nano was applied to serum miRNA profiling for Prostate cancer (PCa) stratification, and machine learning-assisted analysis enabled accurate discrimination among healthy donors (HD), nonmetastatic PCa patients (nPCa), and metastatic PCa patients (mPCa). This work provides a general regenerative nanozyme transduction framework that integrates target recycling, protected catalytic amplification, and multiplexed molecular recognition for liquid biopsy-based cancer stratification.

## 2 Results and Discussion

### 2.1 Principle of TC-DSN-Nano for miRNA Detection

Unlike conventional nucleic acid amplification strategies, TC-DSN-Nano converts miRNA recognition directly into amplified electrochemical outputs through a regenerative transduction mechanism coupled with nanozyme catalysis. As illustrated in Scheme 1, the transducer is constructed by immobilizing liposome-encapsulated gold nanozymes on streptavidin-modified magnetic beads (SA-MBs) through a sequence-programmable DNA linker. Specifically, the DNA linker is dual-functionalized with biotin and cholesterol at its two termini, where the biotin moiety specifically binds to streptavidin on the magnetic bead surface, while the cholesterol moiety anchors the linker into the liposomal lipid bilayer, thereby bridging SA-MB and LA into an integrated transduction assembly. This architecture integrates target recognition, magnetic separation, and signal generation within a single platform. The MBs provide efficient enrichment and purification, whereas the DNA linker serves as both the target-recognition module and the cleavable transduction element. The liposomal carrier serves as a reservoir for a large payload of AuNP nanozymes (liposome@AuNPs, shortened as LA), enabling substantial signal output from each recognition event. Beyond serving as a high-capacity carrier, the liposomal architecture plays an important role in preserving AuNP nanozyme activity during transducer fabrication. The traditional way to directly immobilize AuNP nanozymes on solid supports typically requires extensive surface modification and exposes catalytic sites to biomolecules adsorption, which would reduce catalytic efficiency through active-site blockage and steric hindrance. In contrast, the present design spatially separates the recognition and catalytic modules. The DNA linker, streptavidin, and MBs are localized outside the lipid membrane, whereas the AuNP nanozymes stay confined within the liposomal interior. This compartmentalized configuration minimizes undesired surface passivation and preserves the intrinsic catalytic activity of the AuNP nanozymes throughout probe assembly and target recognition.

Upon target introduction, the miRNA hybridizes with the complementary DNA linker to form a DNA-miRNA duplex. DSN selectively recognizes and cleaves the DNA strand within the heteroduplex, resulting in the detachment of LA from the MBs. Because the miRNA remains intact after cleavage, it is continuously recycled to initiate successive cleavage events. This catalytic recycling process enables a regenerative signal transduction pathway, in which a single miRNA molecule induces the release of multiple LA, thereby translating low-abundance molecular recognition events into amplified nanoscale outputs. For signal readout, the released LA particles are lysed using Triton X-100 to liberate the encapsulated AuNPs nanozymes. As AuNPs owned strong peroxidase (POD)-like activity, the released AuNPs would catalyze the oxidation of 3,3’,5,5’-tetramethylbenzidine (TMB) in the presence of H_2_O_2_, generating amplified electrochemical signals proportional to the amount of target miRNA. By coupling DSN-mediated target recycling with nanozyme-catalyzed electrochemical amplification, TC-DSN-Nano achieves efficient signal multiplication without enzymatic nucleic acid amplification.

In contrast to PCR-based approaches that rely on reverse transcription and exponential amplification, TC-DSN-Nano establishes a direct molecular-to-electrochemical conversion pathway. The amplification process originates from regenerative target turnover and catalytic signal generation rather than nucleic acid replication, simplifying the analytical workflow while maintaining high sensitivity and reliability. Furthermore, owing to the modular design of the DNA linker, the platform can be readily adapted to different miRNA biomarkers through straightforward sequence reprogramming, providing a versatile framework for multiplexed miRNA detection.

**Scheme 1.**
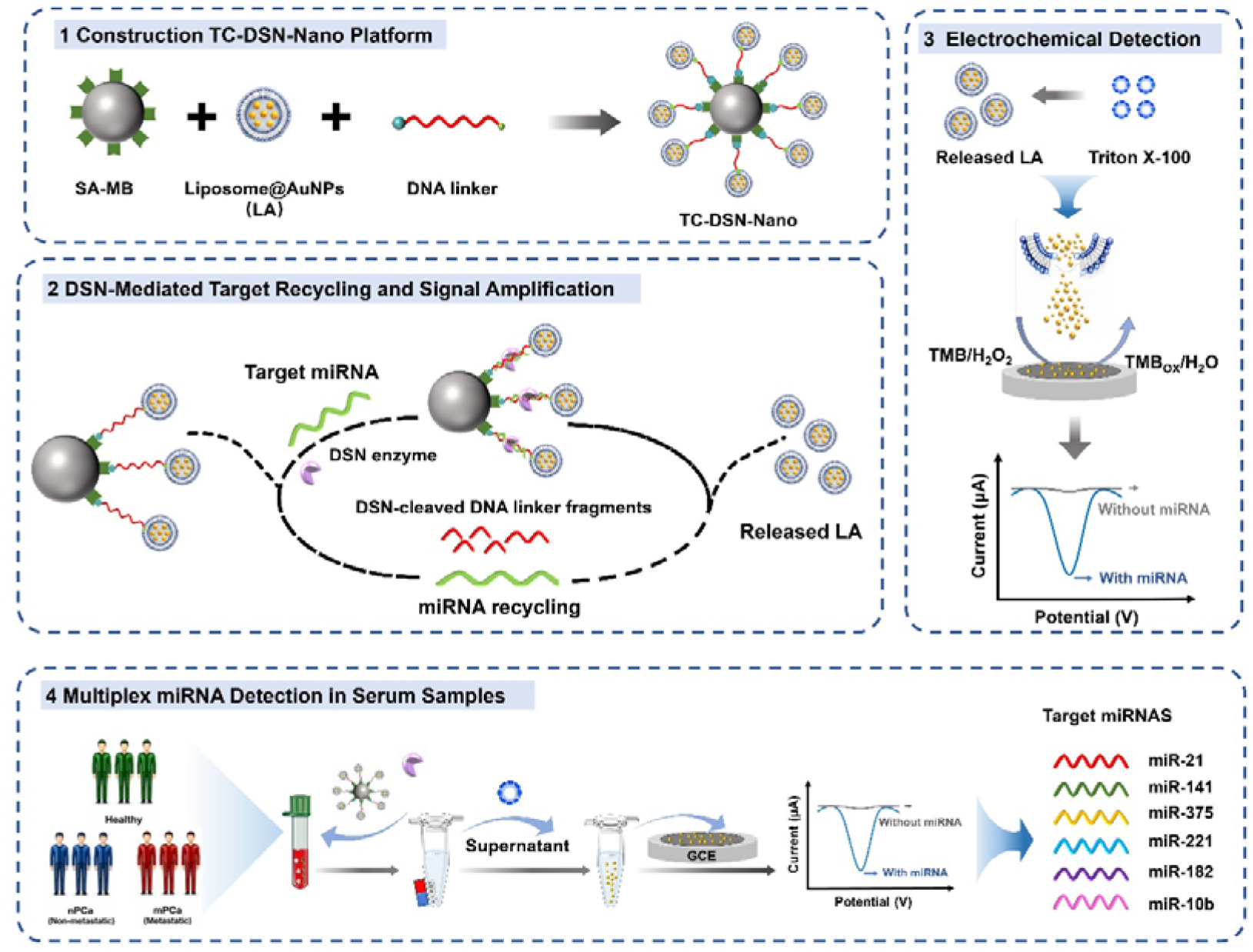
Schematic illustration of TC-DSN-Nano assay for circulating miRNAs profiling. Target miRNA initiates DSN-mediated cleavage of DNA linkers and regenerative release of LA. The liberated AuNPs catalyze the H_2_O_2_/TMB reaction, producing amplified electrochemical signals for sensitive miRNA quantification.

### 2.2 Preparation and Characterization of Liposome@AuNPs

The LA were prepared according to the previously-reported strategy.^36^ AuNPs were formed in situ through diffusion of Au precursor complexes across lipid bilayers, followed by intraliposomal reduction and confined nucleation. As shown in Figure S1 and Figure 1A, representative transmission electron microscopic (TEM) images of the LA clearly show lots of AuNPs were surrounded by a thin lipid shell with a thickness of 3.268 nm, confirming nanoparticle formation within the liposomal cavity. Dynamic light scattering analysis showed a slight increase in hydrodynamic diameter following AuNP formation inside the liposomes (Figure 1B). Meanwhile, the zeta potential of blank liposomes and LA showed shift after AuNP encapsulation, suggesting that the incorporation of AuNPs altered the interfacial charge distribution and surface environment of the lipid vesicles (Figure 1C). However, the unchanged liposomal morphology and the absence of obvious surface-deposited nanoparticles in TEM images indicate that AuNPs were primarily encapsulated within the liposomal cavity rather than attached to the outer membrane. The particle size and surface charge of LA remained stable during 7-day storage (Figure S2), demonstrating good colloidal stability of the LA system. UV-vis spectrum further confirmed AuNP formation, as evidenced by the characteristic plasmon resonance band centered at approximately 520 nm (Figure 1D).

**Figure 1.**
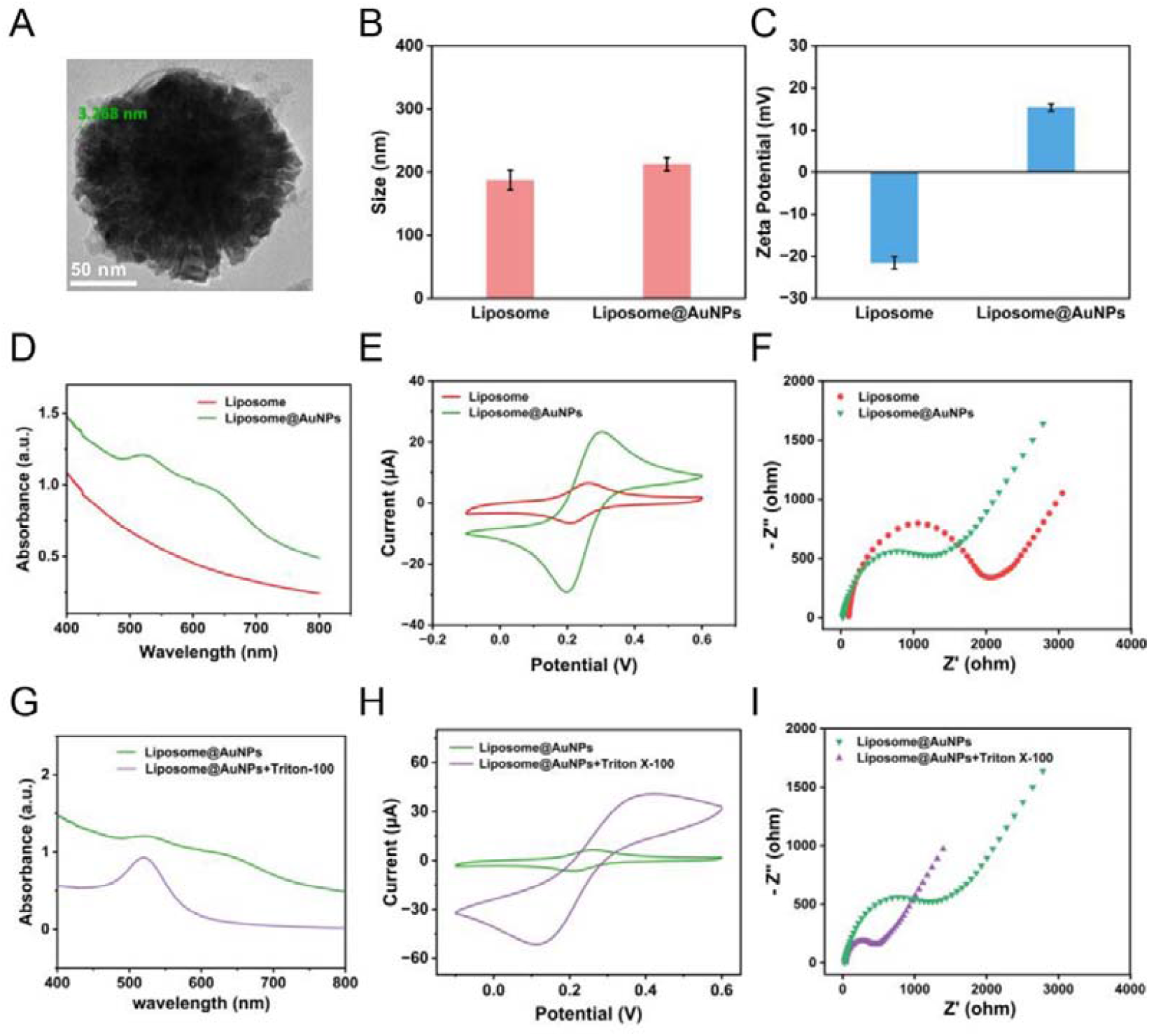
Characterization of LA. (A) TEM image of the LA. (B) Hydrodynamic diameters of liposome and LA. (C) Zeta potential measurement of liposome and LA. (D) UV-vis spectra of liposomes and LA. (E) CV and EIS (F) of LA measurement of liposome and LA on work electrode. (G) UV-vis absorption spectra, (H) CV curves and (I) EIS analysis of LA before and after membrane disruption.

The electrochemical characteristics of LA were subsequently investigated. Cyclic voltammetry (CV) and electrochemical impedance spectroscopy (EIS) measurements revealed slightly enhanced conductivity and electron transfer kinetics compared with empty liposomes due to the presence of internal AuNPs (Figure 1E, F). Nevertheless, the lipid bilayer remained an effective insulating barrier that restricted direct electrical communication between the encapsulated AuNPs and the electrode surface. This feature is advantageous for minimizing background signals prior to target-triggered activation.

To explore the release behavior of AuNPs from LA, 1% Triton X-100 was employed as a membrane-disrupting agent to trigger liposomal disintegration, and the release kinetics of the encapsulated AuNPs were systematically evaluated by monitoring the effects of incubation temperature and time. The increase in absorbance at 520 nm was used as an indicator of AuNPs exposure and release. Under the investigated conditions, incubation at 60 °C for 30 min resulted in a plateau in the signal, suggesting efficient disruption of the liposomal membrane and near-complete release of the encapsulated AuNPs (Figure S3). The release process was further investigated to verify the efficiency and kinetics of membrane disruption. The power output curve for Triton X-100 in time-dependent release studies further confirmed efficient disruption of the lipid membrane and rapid liberation of encapsulated AuNPs (Figure S4).

To further verify AuNP release and its associated electrochemical consequences, multiple complementary characterization methods were employed. UV-vis spectra exhibited a significant increase in absorbance after Triton X-100 treatment (Figure 1G), indicating LA lysis and AuNPs release. Consistently, CV measurements demonstrated a significant increase in current response following membrane disruption (Figure 1H), while EIS revealed a substantial decrease in charge-transfer resistance (Figure 1I), indicating marked interfacial electron transfer after AuNPs release. These results demonstrate that Triton X-100-induced membrane disruption efficiently exposes the encapsulated AuNPs and restores their electrochemical accessibility.

Beyond facilitating AuNP release, the liposomal membrane also plays an important protective role during transducer construction. Specifically, because the catalytic AuNPs are shielded from direct contact with DNA linkers, proteins, and magnetic supports during transducer construction, their catalytic surfaces remain largely unperturbed. Upon liposome disruption, these intact AuNPs are released and become fully accessible for catalysis, thereby maximizing signal output. This protective effect is further supported by the marked enhancement in electrochemical response following Triton X-100 treatment, suggesting that the liposomal compartment effectively preserves the catalytic activity of the encapsulated AuNPs prior to activation. Therefore, the lipid membrane effectively isolates the electroactive AuNPs before activation while enabling rapid signal amplification upon lysis. In this regard, the liposome functions not only as a signal reservoir but also as a protective nanoreactor that preserves AuNP nanozyme activity while enabling efficient target-to-signal conversion.

The electrochemical behavior of the released AuNPs was further examined using CV at different scan rates. The peak current increased progressively with increasing scan rates (Figure S5A). A linear relationship was observed between the anodic and cathodic peak currents and the square root of scan rate (Figure S5B), indicating a diffusion-controlled electrochemical process. These results demonstrate efficient electron-transfer kinetics of the released AuNPs introduced on the electrode surface and support their suitability as catalytic signal reporters for electrochemical biosensing.

### 2.3 Engineering of a DSN-Mediated Transducer (TC-DSN-Nano) for miRNA Assay

To demonstrate the DSN-mediated cleavage mechanism, miRNA-DNA heteroduplexes were first generated by hybridizing the target miRNA with its complementary DNA linker. As shown in Figure 2A, the intact miRNA-DNA heteroduplex exhibited a distinct band corresponding to the duplex structure in the absence of DSN. Upon DSN treatment, the intensity of the heteroduplex band gradually decreased with increasing incubation time, accompanied by the appearance of shorter DNA fragments, indicating the efficient cleavage of the DNA strand within the RNA-DNA heteroduplex. In contrast, the miRNA strand remained resistant to DSN-mediated digestion and was was regenerated after cleavage, allowing it to participate in subsequent rounds of hybridization and DNA cleavage. These results demonstrate that DSN can selectively cleave the DNA linker in the miRNA-DNA heteroduplex while preserving the target miRNA, thereby enabling miRNA regeneration, and establishing the basis for catalytic signal amplification in the proposed transduction system.

**Figure 2.**
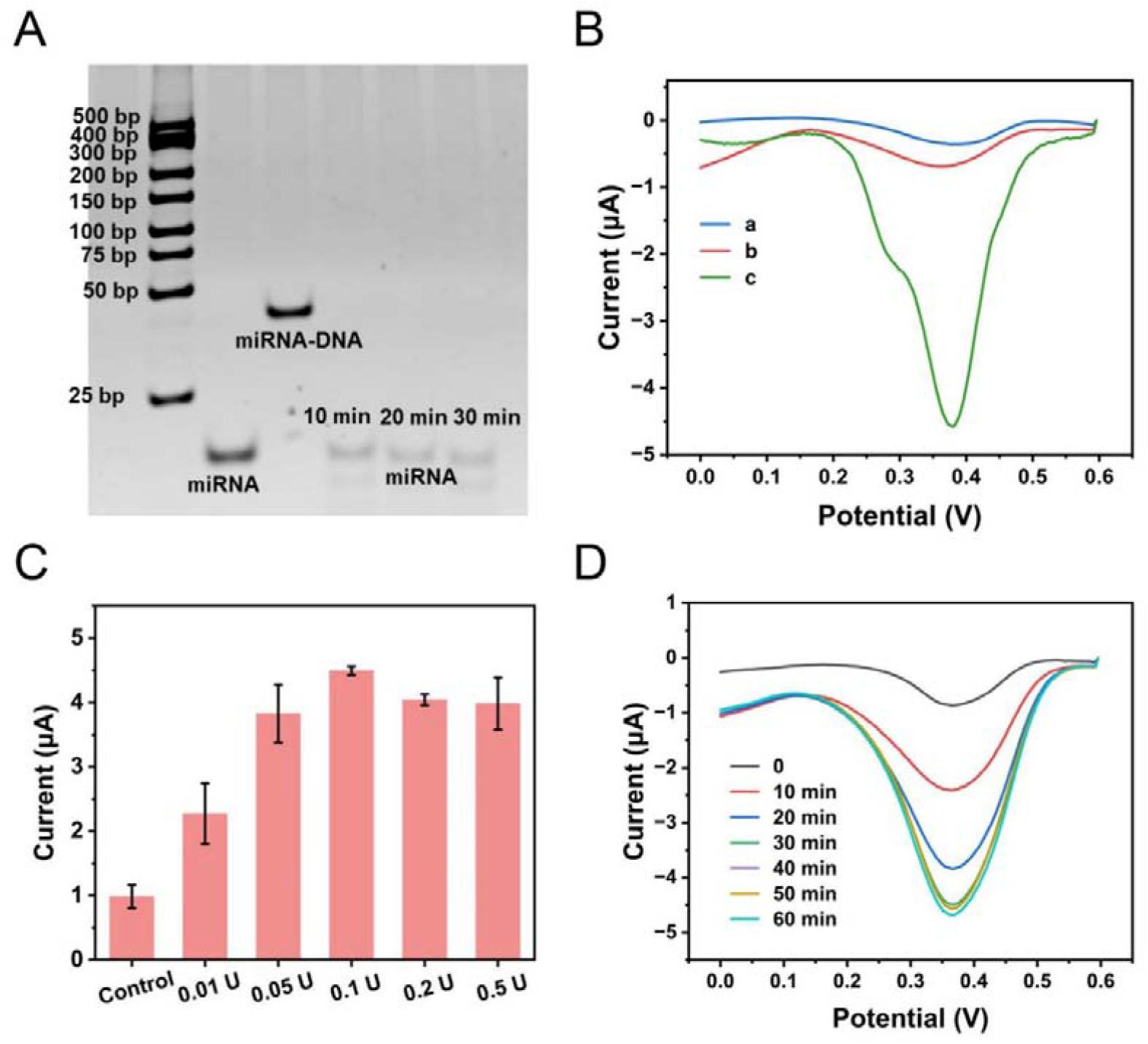
Establishment of the DSN-mediated regenerative transduction mechanism. (A) Gel electrophoresis analysis of miRNA-DNA heteroduplexes cleavage by DSN enzyme. (B) DPV responses of TC-DSN-Nano under different activation conditions: the absence of the target miRNAs (a), without DSN (b), and with both miRNA-141 and DSN. (C) Electrochemical signal on adding increasing amounts of DSN enzyme. (D) Time-dependent electrochemical response of released LA. The concentration of miRNA was 10 pM. Error bars are the standard deviation of three repetitive experiments.

The functionality of the TC-DSN-Nano platform was subsequently evaluated by electrochemical measurements. As shown in Figure 2B, negligible current responses were observed in the absence of target miRNA or DSN (curves a, b), indicating that spontaneous release of LA was effectively suppressed. Under these conditions, the DNA linker remained intact and the transducer retained its assembled structure on the MBs. In contrast, simultaneous introduction of both target miRNA and DSN produced a pronounced increase in current response (Figure 2B, curve c), confirming efficient activation of the transducer. The enhanced signal originated from DSN-mediated cleavage of the DNA linker, followed by regenerative target recycling and release of LA. Subsequent LA lysis activates AuNP nanozymes, resulting in improved electrochemical accessibility and amplified signal output.

Next, taking miR-141 as an example, we optimized the amount of DSN enzyme. An optimal electrochemical difference in the presence and absence of miRNA was observed when DSN was 0.1 U in a 30 μL reaction mixture (Figure 2C). Hence, 0.1 U of DSN was used throughout subsequent experiments. To further investigate the release behavior of the transduction process, the LA were monitored by measuring the electrochemical response of the supernatant at different target concentrations. As shown in Figure 2D, a time-dependent increase in electrochemical signal was observed, indicating continuous release of LA driven by DSN-mediated cleavage. A peak current increased immediately and then tended to stabilize after 30 min. Notably, the magnitude of signal enhancement strongly depended on target concentration. Samples containing higher concentrations of miRNA generated a significantly stronger electrochemical response throughout the reaction process, whereas lower concentrations generated slower signal accumulation and reduced signal intensity. This concentration-dependent release behavior reflects the increased frequency of miRNA-triggered cleavage and target recycling events at higher target abundance. These results demonstrate that TC-DSN-Nano system efficiently transduces miRNA concentration into a time-dependent electrochemical output, proving a robust basis for quantitative miRNA analysis.

### 2.4 Multiple miRNA Analysis by Regenerative Signal Transduction

The analytical performance of TC-DSN-Nano was systematically evaluated using three PCa-associated miRNAs, including miR-141, miR-21, and miR-10b, as representative targets. As shown in Figure 3A-C, the electrochemical response progressively strengthened with increasing target concentration for all the studied three miRNAs over the concentration range of 0-500 pM, indicating efficient signal transduction driven by efficient coupling between DSN-mediated target recycling and nanozyme-based electrochemical amplification. Quantitative analysis revealed a linear relationship between the peak current change and the logarithm of target concentration for all three miRNAs (Figure 3D-F) over the tested concentration ranges. Based on the corresponding calibration curves, the limits of detection (LODs) were calculated to be 18.4 fM for miR-141, 19.8 fM for miR-21, and 20.7 fM for miR-10b, respectively, demonstrating the high sensitivity and multiplexed detection capability of TC-DSN-Nano for miRNA analysis. The consistent analytical performance obtained across different targets highlights the sequence-programmable nature of the platform, where target recognition is solely determined by the DNA linker sequence without altering the signal transduction mechanism. This modular design enables straightforward adaptation to diverse miRNA biomarkers while maintaining a uniform sensing workflow.

**Figure 3.**
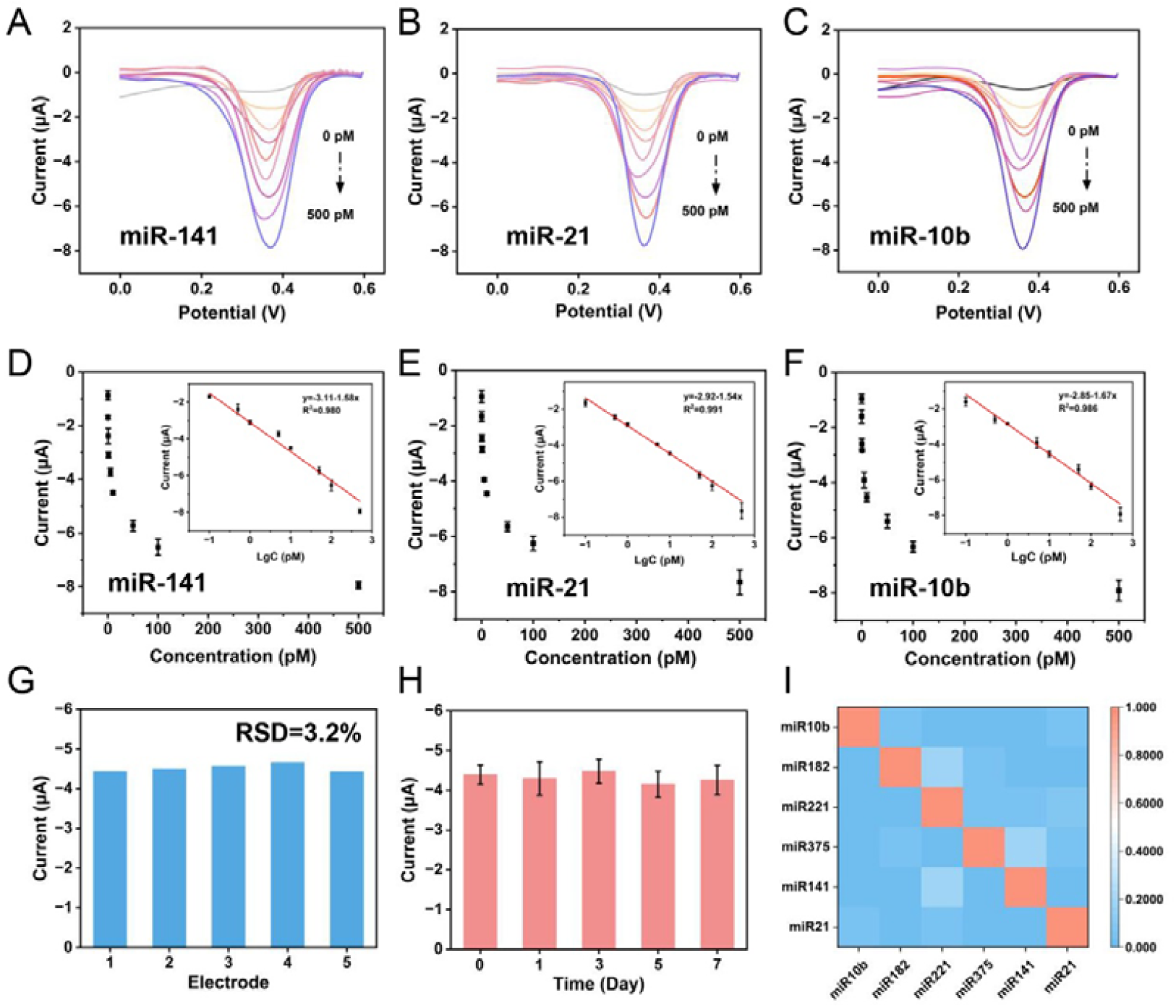
Analytical performance of the TC-DSN-Nano electrochemical sensing platform. (A-C) DPV responses toward different concentrations of miR-141, miR-21, and miR-10b, respectively. (D-F) Calibration curves for quantitative miRNA analysis. Insets show the linear relationships between the peak current and the logarithm of miRNA concentration. (G) Reproducibility of the sensing platform using independently fabricated electrodes. (H) Storage stability over 7 days. (I) Selectivity test representing correlation heatmap analysis of target and non-target miRNAs. Error bars represent the standard deviations (n = 3).

The reproducibility and operational stability of the platform were subsequently evaluated to assess the reliability of the sensing architecture. As shown in Figure 3G, five independently fabricated sensors produced highly consistent responses toward the same concentration of target miRNA, yielding a relative standard deviation (RSD) of 3.2%. The low variation indicates reliable probe fabrication and robust signal transduction. Storage stability was further evaluated over a period of 7 days (Figure 3H). Only minor fluctuations in signal intensity were observed during the testing period, suggesting that the transducer architecture remained structurally and functionally stable under the storage conditions employed.

The sequence specificity of TC-DSN-Nano was further examined using target and non-target miRNAs. As illustrated in the correlation heatmap (Figure 3I), each sensing probe generated the strongest response toward its perfectly matched target, whereas only weak responses were observed from mismatched or non-complementary sequences. This high selectivity originates from the stringent hybridization requirement between the target miRNA and DNA linker, which governs subsequent DSN-mediated cleavage and signal generation. Because efficient transduction can only occur after formation of a matched miRNA-DNA duplex, nonspecific activation is effectively suppressed. These results demonstrate that the TC-DSN-Nano platform provides broad target adaptability, quantitative response, high reproducibility, operational stability, and sequence-specific recognition within a unified sensing framework.

The observed analytical performance can be attributed to the synergistic integration of DSN-mediated miRNA recycling and liposome-protected nanozyme amplification. Upon target recognition, DSN-mediated cleavage continuously regenerates the target miRNA, while the controlled release of intact AuNP nanozymes enables efficient catalytic signal amplification, thereby generating an electrochemical response proportional to the target concentration. By coupling DSN-mediated target recycling with liposome-protected and controllably released nanozyme amplification, TC-DSN-Nano achieves efficient signal multiplication without enzymatic nucleic acid amplification.

### 2.5 Serum miRNA Profiling for Prostate Cancer Stratification

To evaluate the clinical applicability of the TC-DSN-Nano, serum samples were collected from an age-matched cohort comprising HD (*n* = 12), nPCa (*n* = 20), and mPCa (*n* = 20). Six circulating miRNAs associated with PCa progression, including miR-141, miR-21, miR-182, miR-375, miR-221 and miR-10b, were selected to establish a multi-marker serum signature.^9, 37-40^ The distribution of electrochemical signals for the six miRNAs is summarized in Figure 4A. The expression profiles of the six miRNAs are summarized in the heatmap (Figure 4B). Most of these miRNAs displayed elevated expression from HD to nPCa and further to mPCa. Although the magnitude of change varied among individual biomarkers, the collective expression pattern revealed a clear disease-associated trajectory, suggesting that integrated miRNA profiling captures molecular alterations accompanying PCa progression more effectively than any single biomarker.

**Figure 4.**
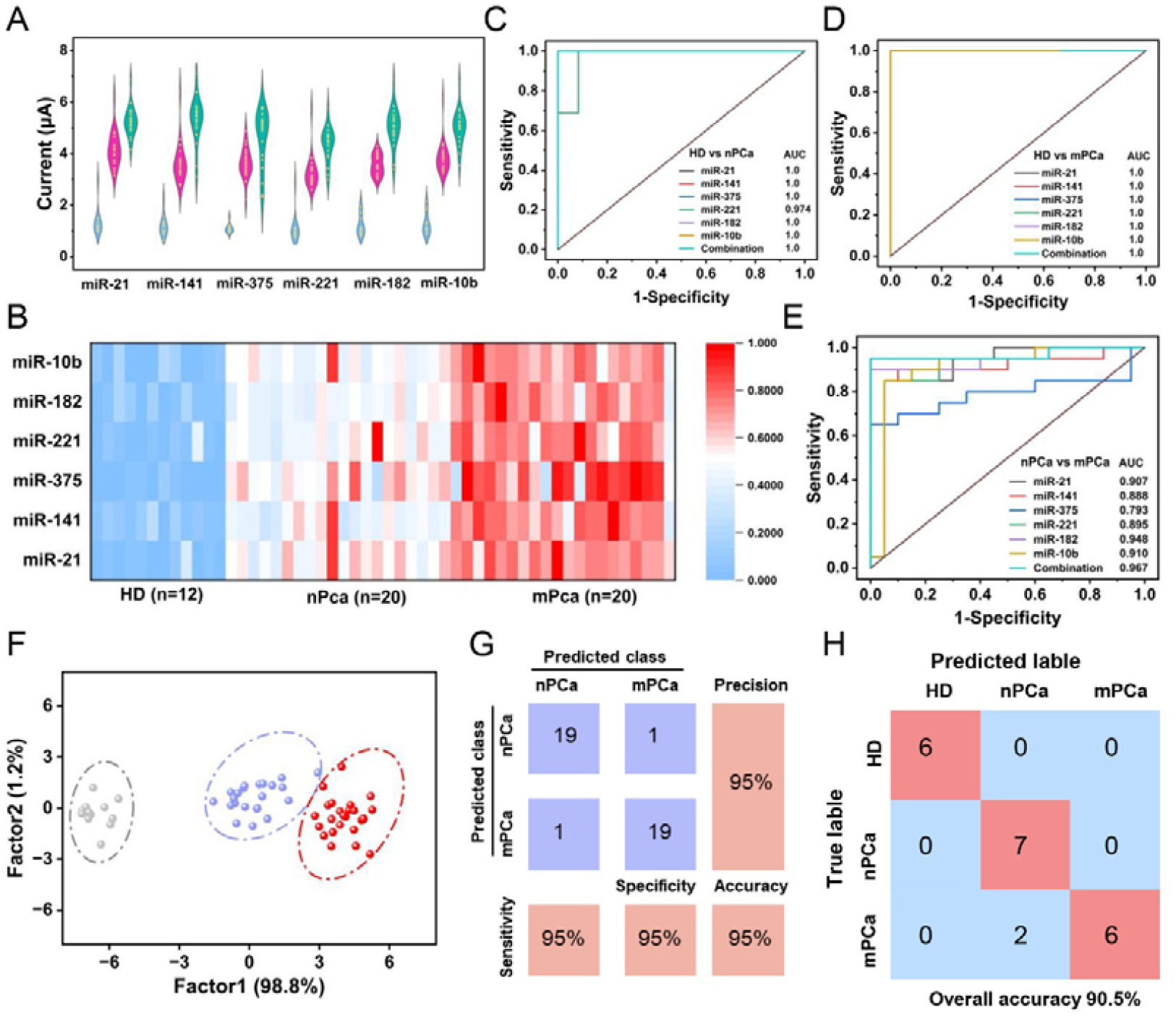
Clinical evaluation of the TC-DSN-Nano for serum miRNA profiling and PCa stratification. (A) Electrochemical responses of the six target miRNAs in HD (blue color), nPCa (red color), and mPCa (green color) serum samples. (B) Heatmap of six circulating miRNAs measured in serum samples from HD (n=12), nPCa (n=20), and mPCa (n=20). (C-E) ROC curves for distinguishing (C) HD from nPCa, (D) HD from mPCa, and (E) nPCa from mPCa using individual miRNAs and the combined six-miRNA panel. (F) and (G) LDA score plot based on the six-miRNA signature. (H) Confusion matrix of the XGBoost classification model based on six-miRNA signature.

The diagnostic performance of individual miRNAs and the combined biomarker panel was further evaluated by using receiver operating characteristic (ROC) analysis (Figure 4C-E). Across all three comparison scenarios (HD vs. nPCa, HD vs. mPCa, and nPCa vs. mPCa), the combined six-miRNA panel consistently achieved AUC values comparable to or higher than those of individual biomarkers. The advantage of the combined signature was particularly evident in discriminating mPCa from nPCa, demonstrating that multiplexed miRNA profiling provides more informative molecular signatures for disease stratification than individual biomarkers.

To determine whether the six-miRNA signature could support disease stratification, multivariate analysis was performed using the electrochemical profiling data. As shown in Figure 4F and Figure S6, linear discriminant analysis (LDA) and principal component analysis (PCA) were used to enhance the performance of predictive models. Projection of the samples onto the first two discriminant components resulted in three relatively distinct clusters corresponding to HD, nPCa, and mPCa. Clear separation was observed between HD and PCa, while metastatic and non-metastatic cases also formed distinguishable groups with only limited overlap. These results indicate that the multidimensional molecular information obtained from TC-DSN-Nano preserves biologically relevant differences associated with disease stage and metastatic status.

To further evaluate the predictive value of the multiplexed miRNA signature, an XGBoost model was established using the electrochemical responses of all six miRNAs. As shown in Figure 4G, the confusion matrix yielded an overall classification accuracy of 90.5%, with only a small number of misclassified samples. Most HD, nPCa, and mPCa were correctly assigned to their respective groups, indicating that the multidimensional molecular information generated by TC-DSN-Nano effectively capture disease-associated differences and support reliable clinical classification. Taken together, these results demonstrate that multiplexed profiling of circulating miRNAs using the TC-DSN-Nano enables substantially greater discriminatory power than individual biomarkers alone. The integrated molecular signatures facilitate accurate PCa detection and metastatic stratification, highlighting the potential of TC-DSN-Nano as a minimally invasive tool for precision cancer diagnostics.

## 4. Conclusion

In summary, we developed a TC-DSN-Nano platform that establishes a regenerative nanozyme transduction strategy for multiplexed circulating miRNA profiling. By integrating DSN-mediated target recycling with a liposome-protected AuNP nanozyme reporter, the platform spatially separates molecular recognition from signal generation and enables direct conversion of miRNA-triggered events into amplified electrochemical outputs without conventional nucleic acid amplification. The liposomal compartment preserves nanozyme catalytic activity during transducer assembly, while target-activated release of AuNPs provides efficient signal amplification for sensitive and programmable miRNA quantification. Coupled with multivariate analysis and machine learning-assisted classification, TC-DSN-Nano-derived multiplexed miRNA signatures enabled accurate discrimination among HD, nPCa, and mPCa patients, demonstrating the value of integrating molecular profiling with data-driven disease stratification. The modular design allows facile adaptation to diverse nucleic acid targets through sequence reprogramming, providing a generalizable framework for liquid biopsy applications and multiplex analysis of low-abundance biomarkers in complex biological systems.

## Associated Content

### Data Availability Statement

The research data presented in this study are available on request from the corresponding author.

### Supporting Information

The Supporting Information is available free of charge at http://pubs.acs.org/xxxxx.

Materials and reagents, Synthesis of liposome and Liposome@AuNPs, miRNA-responsive transducer assembly, Regenerative signal transduction, The kinetics of LA release, Electrochemical measurements (PDF).

## Acknowledgements

This work is supported by the National Natural Science Foundation of China (Grant Nos. 22174049, 22574059), the Natural Science Foundation of Hubei Province of China (No. 2021CFB335). The authors also thank the Analytical and Testing Center of HUST and the Medical Subcenter of HUST Analytical & Testing Center for material characterization and data acquisition.

